# It takes a dimer to tango: Oligomeric small heat-shock proteins dissociate to capture substrate

**DOI:** 10.1101/369454

**Authors:** Indu Santhanagopalan, Matteo T. Degiacomi, Dale A. Shepherd, Georg K.A. Hochberg, Justin L.P. Benesch, Elizabeth Vierling

## Abstract

Small heat-shock proteins (sHsps) are ubiquitous molecular chaperones, and their mutations or altered expression are linked to multiple human disease states. sHsp monomers assemble into large oligomers with dimeric substructure, and the dynamics of sHsp oligomers has led to major questions about the form that captures substrate, a critical aspect of their mechanism of action. We show that substructural dimers of plant dodecameric sHsps, Ta16.9 and homologous Ps18.1, are functional units in the initial encounter with unfolding substrate. We introduced inter-polypeptide disulfide bonds at the two dodecameric interfaces, dimeric and non-dimeric, to restrict how their assemblies can dissociate. When disulfide bonded at the non-dimeric interface, mutants of Ta16.9 and Ps18.1 (Ta_CT-ACD_ and Ps_CT-ACD_) were inactive, but when reduced had wild-type-like chaperone activity, demonstrating that dissociation at non-dimeric interfaces is essential for activity. In addition, the size of the Ta_CT-ACD_ and Ps_CT-ACD_ covalent unit defined a new tetrahedral geometry for these sHsps, different than the Ta16.9 x-ray structure. Importantly, oxidized Ta_dimer_ (disulfide bonded at the dimeric interface) showed greatly enhanced ability to protect substrate, indicating that strengthening the dimeric interface increases chaperone efficiency. Size and secondary structure changes with temperature revealed that folded sHsp dimers interact with substrate, and support dimer stability as a determinant of chaperone efficiency. These data yield a model in which sHsp dimers capture substrate prior to assembly into larger, heterogeneous sHSP-substrate complexes for subsequent substrate refolding or degradation, and suggest that tuning the strength of the dimer interface can be used to engineer sHsp chaperone efficiency.

## INTRODUCTION

Small heat-shock proteins (sHsps) are a class of ATP-independent chaperones, expressed across all kingdoms of life, that are proposed to act as a cell’s ‘first-responders’ under stress conditions (1-6). Mutations or mis-expression of specific human sHsps are associated with myopathies, neuropathies and cancers (2, 7-9). The canonical function of sHsps is that they capture substrate proteins that are partially denatured by heat or other stresses in large complexes, which are acted upon by ATP-dependent molecular machines to promote either substrate refolding or degradation (2, 10).

sHsps are characterized by a ~90 amino acid, β-sheet rich α-crystallin domain (ACD, or Hsp20 domain, PF00011), flanked by sequences of variable length and composition (N-terminal sequence or NT; C-terminal sequence or CT) (11, 12). The majority form large oligomers containing 12 to >40 units, with a dimeric substructure (2, 3, 5), although there are a few dimeric sHsps, some of which are active chaperones (5, 13-15). While vertebrate sHsps principally assemble as polydisperse oligomers, some bacterial, yeast and plant sHsps are monodisperse. In all oligomeric forms, sHsp dimers interact through a conserved interface between an I-X-I motif in the CT and a groove formed by β4 and β8 in the ACD of a monomer in another dimer (hereafter the CT-ACD interface) (2, 5, 16, 17). Notably, sHsp oligomers are also highly dynamic, exchanging dimers (and monomers) on a timescale of minutes, as fast or faster than their rate of substrate capture.

The large size, dynamic nature and inherent polydispersity of most sHsps have resulted in few high resolution structures (17-22) and differing hypotheses on the sHsp mode of action. Interaction with substrates has been proposed to require either conformational changes in the oligomer or dissociation of oligomers into active units (3-5). A mechanism involving structural rearrangements of the oligomers is supported by presence of multiple oligomer conformations (23-25), with a shift in equilibria proposed to activate the sHsp. This model is also supported by experiments indicating that sHsp chaperone activity is unaffected by mutations or cross-linking that alter rates of subunit exchange (26-28). In contrast, multiple studies have measured temperature-dependent subunit exchange of several sHsps, consistent with a model in which sHsp dissociation releases suboligomers that interact with unfolding proteins (29-37) (38). In this model, the oligomeric form sequesters the major substrate binding regions of the sHsp, which is further supported by analysis of crosslinking interactions with substrates (39).

Higher plants express large numbers of sHsps belonging to 11 gene families (40), likely owing to their ‘immobile’ life that exposes them to environmental changes. Understanding sHsp function has implications for engineering plants to withstand temperature changes, as well as for defining their role in multiple human disease states. We have aimed to ascertain the mechanism and substrate binding unit of sHsps through studies of cytosolic class-I sHsps from plants, which have proven to be exceptional models for the study of sHsp chaperone mechanism. Their largely mono-disperse, dodecameric form under ambient conditions (41-43) and the availability of a high resolution structure for wheat (*Triticum aestivum*) sHsp Ta16.9 (PDB 1GME) (17) have made them amenable to functional and mechanistic studies. It has been suggested that these dodecamers, as well as oligomers of other sHsps, are reservoirs of dimers that are released under stress conditions to interact with unfolding substrates (6, 44). In this study, we used Ta16.9 and its close homologue from *Pisum sativum*, Ps18.1 (5, 17, 41), to address models of sHsp substrate interaction. These sHsp dodecamers display temperature-dependent dynamics in subunit exchange studies, with exchange mediated primarily by dimers (33-35, 45). Analysis of stoichiometries of substrate-sHsp complexes for Ps18.1 has shown that there is a bias for an even number of sHsp subunits in sHSP-substrate complexes (44, 46). However, there is no direct evidence for sHsp dissociation being essential for activity or for dimers being important for substrate capture.

To address the mechanism of substrate protection by sHsps, we engineered disulfide bridges that prevent dissociation at either of the two major oligomeric interfaces, dimeric and non-dimeric, in the Ta16.9 and Ps18.1 dodecamers, and tested their activity in protecting a heat-sensitive protein, malate dehydrogenase (MDH). The mutants led us to re-examine the quaternary structures of Ta16.9 and Ps18.1 and support the conclusions that sHsps need to dissociate at specific interfaces to protect substrate, and that free, folded dimers are the active substrate-capturing units.

## RESULTS

### Protein design, expression, and purification

To test the functional importance of sHsp oligomer dynamics, we made cysteine mutant pairs in Ta16.9 and Ps18.1 to prevent dissociation at the non-dimeric or dimeric interfaces. To link the non-dimeric interfaces, cysteine residues were introduced to create a disulfide bridge between the CT of one monomer and β4 of the ACD from a monomer in another dimer (**Fig. 1A**). Of cysteine pairs tested, the mutants E74C V144C Ta16.9 (Ta_CT-ACD_) and E81C V151C Ps18.1 (Ps_CT-ACD_) were soluble and purified as dodecamers (Table S1, **Fig.** S1). To prevent dissociation at the dimeric interface, cysteines were introduced in a non-hydrogen-bonded registered pair between β6 on one monomer and β2 of the partner subunit such that the resulting disulfide-bonded dodecamer would dissociate into six covalent dimers when oxidized (**Fig. 1A**). The mutant K49C W96C Ta16.9 (Ta_dimer_) was soluble and purified as a dodecamer; we were unable to create an analogous, dodecameric Ps_dimer_, and studies on this interface are confined to the Ta_dimer_ protein. This interface, which involves strand swapping between ACDs, is characteristic of plant, bacterial, and yeast sHsps (3, 5, 17-19).

**Figure 1.**
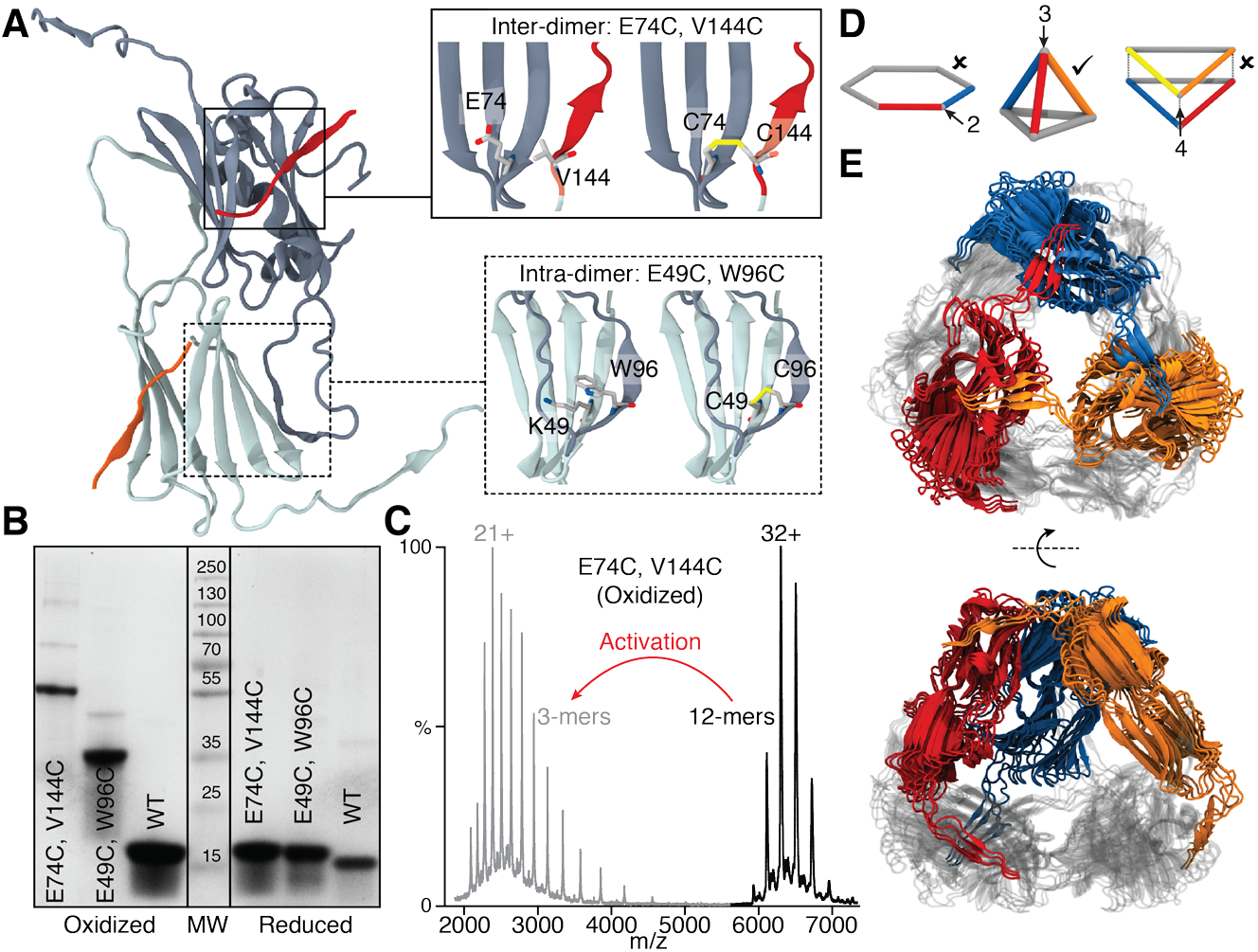
Disulfide bonds introduced within the sHsp dimer or between sHsp dimers, the latter of which defines a new geometry of the sHsp dodecamer. **(A)** Positions of the Cys substitutions linking dimers (Inter-dimer; top inset) with the CTD of one dimer shown in red, and a segment of the β-sandwich of the other dimer in blue. Positions of the Cys substitutions linking monomers within a dimer (Intra-dimer; bottom inset) with segments of one monomer in dark blue and the other monomer in light blue. **(B)** SDS-PAGE separation of the indicated wheat wild type (WT) or mutant proteins either oxidized (left) or reduced (right). **(C)** Mass spectra of Ta E74C V144C (Ta_CT-ACD_) in the oxidized state before (black) and after activation (MS2) of the 32+ ion (gray), showing dissociation to covalent trimers. **(D)** The possible geometries of dodecamers comprising dimers with each edge corresponding to a dimer. The only geometry consistent with a three point linkage is a tetrahedron (middle). **(E)** Tetrahedral models of Ta16.9 based on SAXS and collisional cross section data from ion mobility mass spectrometry (see text and Supplemental figures and methods for details). Three dimers are colored (blue, orange, red) and three are rendered in gray.

The formation of disulfide bonds and absence of free cysteines in all the disulfide mutants after oxidation were confirmed by DTNB assays (Table S2). To ascertain if the desired disulfide bonds were formed, the wheat proteins were examined by SDS-PAGE in the presence or absence of a reducing agent (**Fig. 1B**). The reduced proteins migrated at the expected monomeric size, while without reducing agent Ta_CT-ACD_ and Ta_dimer_ migrated at sizes close to that of a covalent trimer (51 kDa) and covalent dimer (34 kDa), respectively. The apparent size of Ta_CT-ACD_ was surprising, as it was expected to form covalent tetramers (~68 kDa) based on the hexagonal double-ring arrangement in the Ta16.9 crystal structure (PDB:1GME) (17). To verify this observation, we obtained an accurate mass of the non-covalent and covalent units in Ta_CT-ACD_, by using native mass spectrometry (MS) with collisional activation. This confirmed the unexpected result that disulfide linkage at the CT:ACD interface produced covalent trimers rather than tetramers (**Fig. 1C**). The only arrangement featuring dimers connected exclusively in groups of three is a tetrahedron (**Fig. 1D**) (4). Thus, these data can be explained only if the sHsp quaternary arrangement is tetrahedral, in contrast to the double-ring of the Ta16.9 crystal structure (17).

### A new geometry for the sHsp dodecamer

To ensure that Ta_CT-ACD_ had retained the same quaternary structure as the wild-type Ta16.9, we performed native ion mobility mass spectrometry (IM-MS) experiments. Both proteins were dodecameric, and Ta_CT-ACD_ had a very similar collision cross-section (CCS) to the WT, whether oxidized or reduced (Fig. S2), indicating that they are architecturally equivalent. We therefore capitalized on the CCS values to build models for Ta16.9 dodecamers consistent with overall tetrahedral architecture and linkages between ACDs and CTs. An exhaustive search of the roto-translational space, assuming all ACD dimers to be in equivalent environments, and filtering of the models according to the CCS and linkage constraints, resulted in four models that represent the data well (Fig. 1E). All four models have proteins in the tetrahedral architecture with minor differences owing to uncertainty in the CCS measurements.

We did not include the NTs in the modeling due to lack of experimental restraints on this domain, but evidence from the Ta16.9 crystal structure and other sHsps indicates that they reside on the interior of the ACD cage (4). We therefore measured the volume inside the tetrahedral cavity of our models (130395 ± 3060 Å^3^), and calculated the density that would be required in order for 12 NT domains to fit inside. We obtained a density of 0.42 Da/Å^3^, well below the average density of proteins (0.81 Da/Å^3^ (47)), revealing that the cavity is readily able to accommodate 12 NT sequences. This tetrahedral architecture is equivalent to that observed for the sHSP Acr1 from *Mycobacterium tuberculosis* (48). Native MS of Ps18.1_CT-ACD_ (49), and overlapping small angle x-ray scattering (SAXS) profiles for Ta16.9 and Ps18.1 provide evidence that Ps18.1 also has tetrahedral geometry (Fig. S3). Therefore, we conclude that both Ta16.9 and Ps18.1 are primarily tetrahedral dodecamers in solution, and that the cysteine mutations and disulfide bonds do not disrupt this oligomeric architecture.

### Disulfide-bonded sHsps have stabilized secondary structure

We considered that in addition to restricting the mode of dodecamer dissociation, introducing cysteine residues and disulfide bonds could affect the stability of sHsp secondary and quaternary structure, which might alter chaperone activity. Therefore, we first assessed secondary structure of the sHsps at different temperatures by obtaining far-UV circular dichroism (CD) spectra (**Fig. 2A**). Although CD spectra for all the sHsps in the oxidized state and the reduced Ta_dimer_ were recorded, it was not possible to obtain fully reduced Ta_CT-ACD_ or Ps_CT-ACD_ at concentrations of reducing agent compatible with CD (Table S2). Therefore, we created the single cysteine mutants Ta_V144C_ (V144C Ta16.9) and Ps_V151C_ (V151C Ps18.1), to approximate reduced Ta_CT-ACD_ and Ps_CT-ACD_, respectively. The CD spectra for native sHsps are similar to that characteristic of β-sheets, indicating that the CD signal predominantly arises from the ACD β-sandwich (**Fig. 2A**). At 25 °C spectra from all proteins are basically superimposable, but differences are observed already at 45 °C (**Fig. 2A**). Based on the temperature at which the different proteins (at 10 μM) show loss of secondary structure, the relative stability is: oxidized Ps_CT-ACD_ > oxidized Ta_CT-ACD_> oxidized Ta_dimer_ > Ps18.1 ≈ Ps_V151C_ > Ta_V144C_ ≈ reduced Ta_dimer_ ≈ Ta16.9. Clearly, stabilizing the two CT-ACD interfaces or the dimeric interface of each monomer in the oligomer with disulfide bonds, stabilizes sHsp secondary structure.

**Figure 2:**
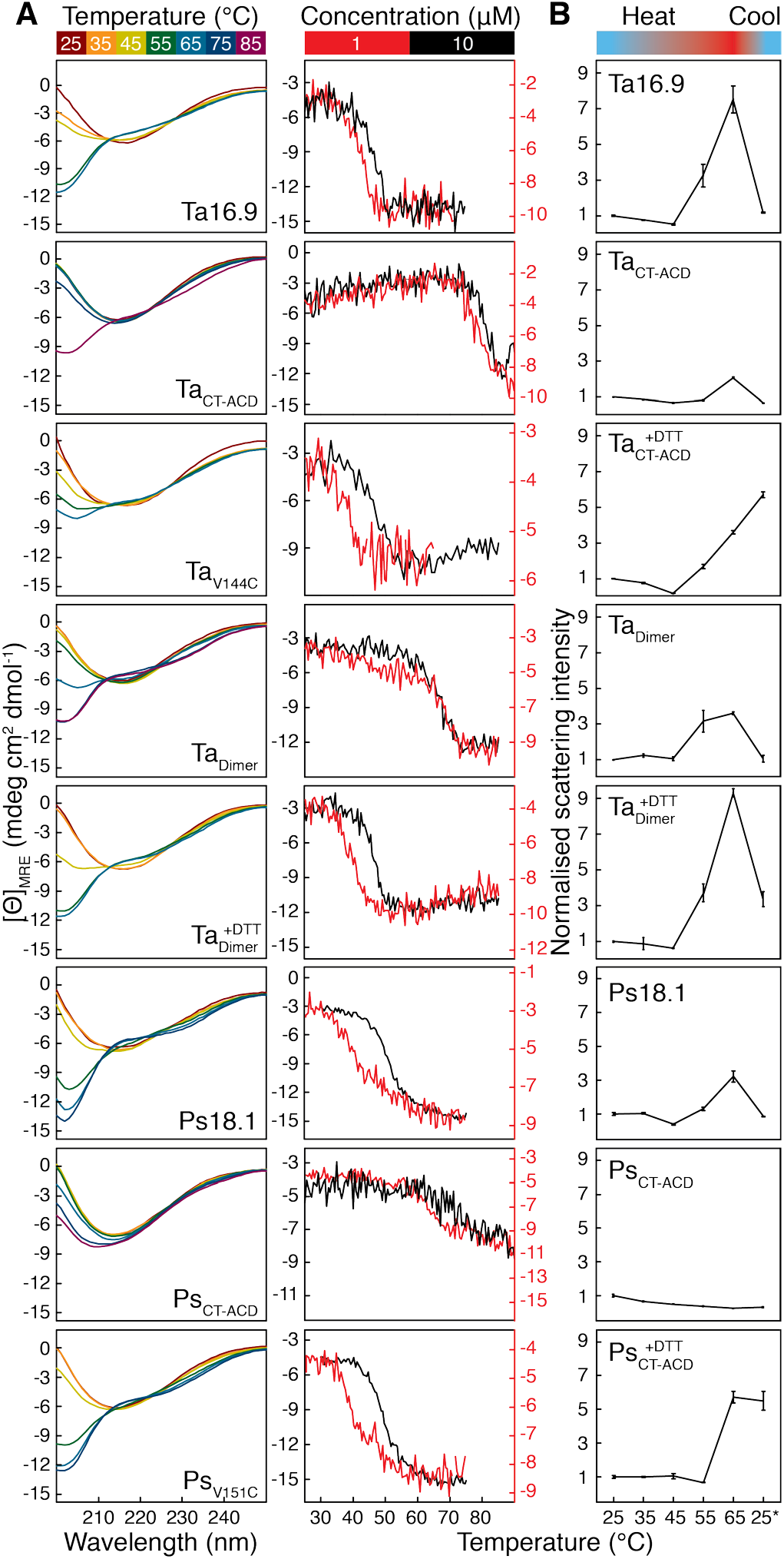
Secondary structure, thermal stability and size of the sHsps. **(A)** Left: Far UV CD spectra for Ta16.9, oxidized Ta_CT-ACD_, Ta_V144C_, oxidized Ta_dimer_, reduced Ta_dimer_, Ps18.1, oxidized Ps_CT-ACD_, and Ps_V151C_ (all 10 μM) collected at temperatures from 25 to 65 °C. Spectra were also collected at 75 and 85 °C for oxidized Ta_CT-ACD_, Ta_dimer_ and Ps_CT-ACD_. Right: Mean Residue Ellipticity (MRE) at 205 nm as a function of temperature for 10 μM (black) and 1 μM (red) sHsp. **(B)** Average values with standard deviations for the total scattering intensities (TSI) of proteins from three experiments carried out at each temperature. Measurements were normalized with respect to 25 °C, which was assigned a value of 1.0.

### Oligomeric state changes with temperature

In order to relate sHsp activity to oligomeric state, we assessed oligomeric transitions in the wild type and mutant proteins at different temperatures. If unfolding is accompanied by changes in sHsp oligomeric state, the unfolding transition temperature will depend on protein concentration. We therefore carried out thermal melts monitored by CD, at 10 and 1 μM sHsp **(Fig. 2A**). For Ta16.9, Ta_V144C_, and reduced Ta_dimer_, cooperative unfolding occurs in the range of 30 to 50 ^o^C, with the transition shifting to a lower temperature at the lower protein concentration. Ps18.1 and Ps_V151C_ behaved similarly, although cooperative unfolding occurred at a higher temperature (40 to 55 °C) (**Fig. 2A**). These data indicate that the oligomeric state of the unfolded and folded species of these proteins are different, and proceed without an intermediate well-folded monomeric state. In contrast, for the oxidized, disulfide-bonded species, Ta_dimer_ and Ta_CT-ACD_, (Ps_CT-ACD_ is highly stabilized, and an unfolded baseline could not be obtained) the temperature range of the unfolding transition, ~60 and 75 ^o^C, respectively, was the same at both protein concentrations (**Fig. 2A**), indicating that their unfolding transition is for the covalent dimer/trimer.

To better understand the oligomeric states involved in substrate protection we examined the temperature dependence of sHsp size by dynamic light scattering (DLS) (**Fig. 2B**, Table S3). When heated to 45 °C, all proteins decreased in total scattering intensity (TSI), consistent with dodecamer dissociation, except oxidized Ta_dimer_ and reduced Ps_CT-ACD_, which showed little change from 25 °C (**Fig. 2B** and Table S3). To relate the TSI to particle sizes, we deconvoluted the DLS correlation curves and broadly classified the particles into dodecameric (7-20 nm diameter), sub-oligomeric (<7 nm diameter) and aggregates (>30 nm diameter) (Fig. S4). At 25 °C, the major species were dodecameric, while at 45 °C, except for reduced Ta_dimer_, the major species for all proteins is smaller. Temperature-dependent formation of both smaller particles and large self-aggregates is consistent with previous studies of sHsps (32, 38, 44, 45). We suggest that the presence of multiple particle sizes explains the unchanged TSI in oxidized Ta_dimer_ and reduced Ps_CT-ACD_, and conclude that all these sHsps undergo dissociation at 45 °C. DLS measurements were also made after cooling the proteins back to 25 °C from 65 ^o^C. Ta16.9, Ps18.1, oxidized Ta_CT-ACD_, Ps_CT-ACD_ and Ta_dimer_ reverted primarily to the dodecameric form, while reduced Ta_CT-ACD_, Ta_dimer_ and Ps_CT-ACD_ were unable to reform dodecamers, but rather formed even larger aggregates. It should be noted that Ta_CT-ACD_, Ps_CT-ACD_ and Ta_dimer_ in their oxidized states, as shown by CD, have native-like secondary structure and are largely folded at 65 ^o^C. The inability of the reduced mutants to revert to dodecamers on cooling was also seen in CD data (Fig. S5), suggesting that the interfaces are critical to refolding.

### Constraining oligomer dissociation alters sHsp chaperone activity

Ta16.9 and Ps18.1 have been well-characterized for their ability to protect the heat-sensitive substrate MDH (17, 41, 50, 51). Therefore, to determine how restricting the ability to dissociate at either the non-dimeric (Ta_CT-ACD_ and Ps_CT-ACD_) or dimeric interfaces (Ta_dimer_) affected sHsp chaperone activity, we assayed the reduced and oxidized sHsps for protection of MDH heated to 45 °C. We first tested to see if the high levels of DTT required to fully reduce the disulfide mutants (so that they would be free to dissociate like wild type) did not interfere with the chaperone activity of the wild type proteins. Both Ta16.9 and Ps18.1 (neither of which have cysteine residues) protected MDH (which has seven cysteine residues) in the presence and absence of reducing agent, as determined by changes in light scattering following heating (**Fig. 3**). As expected, Ps18.1 was more effective in preventing light scattering than Ta16.9, with complete protection of 3 μM MDH at a monomeric molar ratio of sHsp:MDH of 1:1, compared to 6:1 for Ta16.9 (**Fig. 3**)(41, 51). Thus, the wild-type proteins functioned to protect MDH under both reducing and oxidizing conditions.

**Figure 3:**
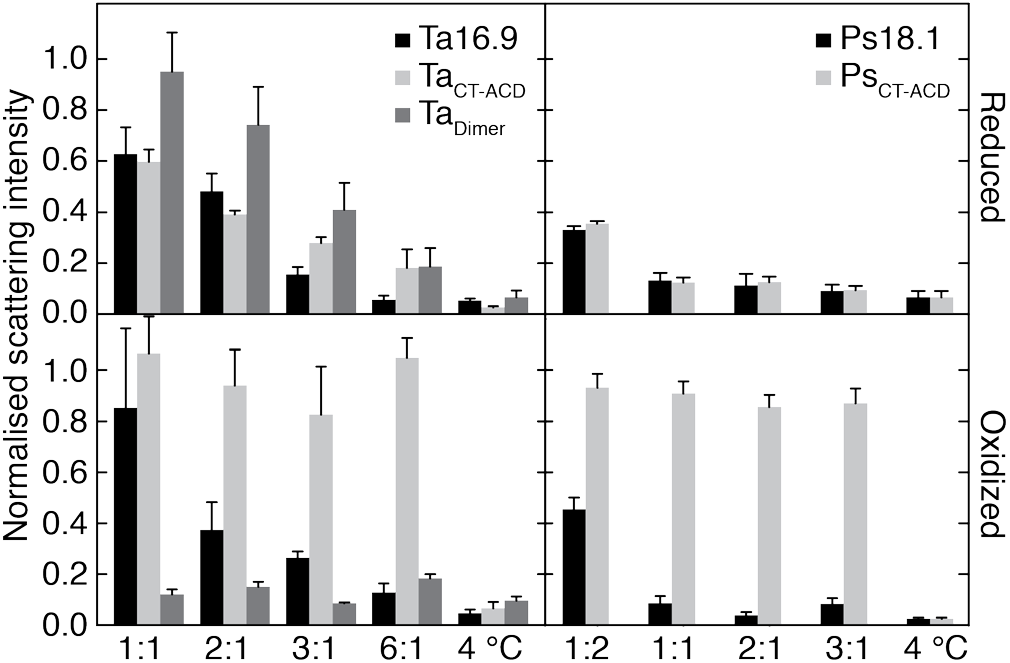
Restricting dodecamer dissociation alters sHsp chaperone activity. Light scattering by aggregated MDH mixed with Ta16.9, Ta_CT-ACD_ or Ta_dimer_ under reducing (Top left) or oxidizing (Bottom left) conditions, and Ps18.1 and Ps_CT-ACD_ under reducing (Top right) or oxidizing (Bottom right) conditions. The sHsp monomer:MDH monomer ratio is indicated on the x-axis. Scattering values were normalized with respect to that for MDH heated alone, which was assigned a value of 1.0. Means from three replicate experiments are plotted with standard error. Unheated MDH (4 ^°^C) served as a control for absence of aggregation.

Comparing the chaperone activity of the reduced and oxidized Ta_CT-ACD_ and Ps_CT-ACD_ revealed that under reducing conditions, both proteins were essentially as effective as their corresponding wild type in preventing MDH light scattering (**Fig. 3**). However, strikingly, under oxidizing conditions, which limits these dodecamers to dissociating into trimers, these sHsps afforded no protection to MDH (**Fig. 3**). Even at a molar ratio of three times that which afforded full protection by Ps18.1, oxidized Ps_CT-ACD_ had no ability to limit MDH light scattering (**Fig. 3**). These data indicate that for the sHsp to protect MDH, it is necessary for the CT:ACD, non-dimeric interfaces to dissociate.

Assays with Ta_dimer_, which can dissociate into dimers under both the reduced and oxidized conditions, yielded dramatically different results. Reduced Ta_dimer_ was somewhat less efficient than Ta16.9 at suppressing MDH aggregation, although still highly effective (**Fig. 3**). In contrast, oxidized Ta_dimer_ protected MDH much more efficiently than Ta16.9, with complete protection at a ratio of Ta_dimer_:MDH of 1:1 (**Fig. 3**). Notably, the CD measurements showed that oxidized Ta_CT-ACD_, Ps_CT-ACD_, and Ta_dimer_ all have highly stable secondary structure, but only oxidized Ta_dimer_ is an effective chaperone. Thus, stable ACD secondary structure alone does not account for the enhanced chaperone activity of oxidized Ta_dimer_, but rather stabilizing the dimeric interface, while still allowing dissociation of dimers from the oligomer, results in a highly effective chaperone.

To examine in more detail how these sHsps interacted with MDH compared to the wild-type proteins, these same samples were analyzed for the presence of MDH-sHsp complexes by size exclusion chromatography (SEC) (**Fig. 4**). As expected, at the sHsp:MDH molar ratios where protection was observed by light scattering, wild-type Ta16.9 and Ps18.1 formed complexes with MDH under reducing or oxidizing conditions, with complexes eluting similarly, ahead of the 670 kDa marker. In contrast, MDH complexes with Ta_CT-ACD_ and Ps_CT-ACD_ were only observed under reducing conditions, when the sHsps are free to dissociate into dimers like wild type; no complexes were formed under oxidizing conditions (**Fig. 4**).

**Figure 4:**
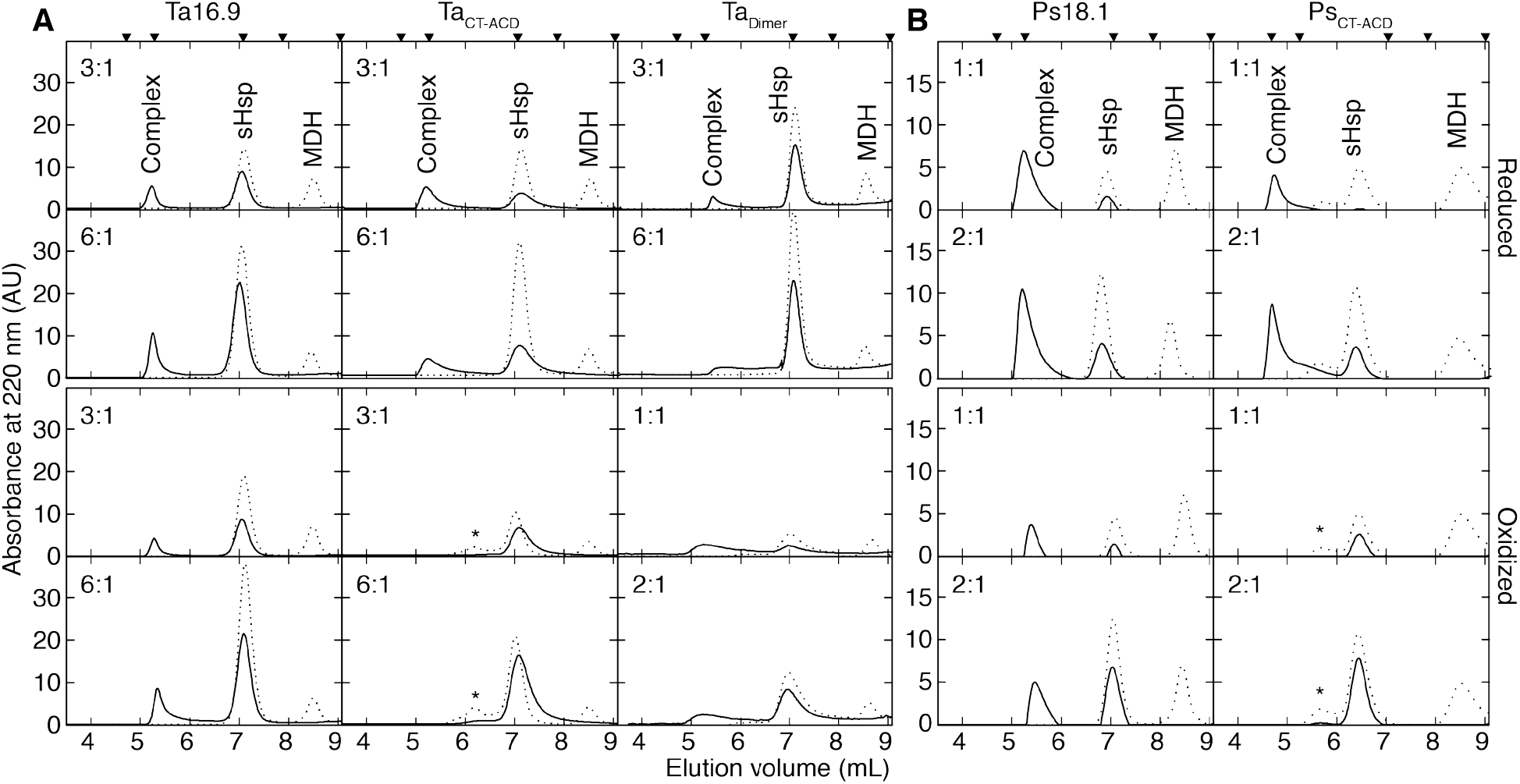
sHsp:MDH complex formation correlates with aggregation protection. SEC profiles were generated on a TSKgel SWxl Ultra SW column for soluble fractions of reaction mixtures of sHsp-MDH under reducing (top 2 rows) or oxidizing conditions (bottom 2 rows) for samples prepared as in Figure 3 with **(A)** Ta16.9 and mutants or **(B)** Ps18.1 and Ps_CT-ACD_. The dashed and solid lines represent unheated and heated reaction mixtures, respectively. The sHsp monomer:MDH monomer molar ratios are indicated at the top right corner of each plot. Calibration standards above the top panels correspond from left to right: Void volume, 670, 158, 44 and 17 kDa. Peaks seen at 7 mL and 8.5 mL correspond to sHsp and MDH, respectively. ‘*’ indicates absorbance from protein with spurious disulfide linkages. Peaks between 4.7-6.5 mL correspond to sHsp-MDH complexes.

Formation of complexes between Ta_dimer_ and MDH also paralleled results of the light scattering assays. The lower efficiency of reduced Ta_dimer_ is reflected in formation of more variable sized Ta_dimer_:MDH complexes compared to Ta16.9:MDH complexes (**Fig. 4**). Conversely, the MDH-sHsp complexes that formed with oxidized Ta_dimer_ include more smaller complexes than those formed with WT Ta16.9, consistent with Ta_dimer_ being a more efficient chaperone (30).

## DISCUSSION

Outstanding questions about the mechanism of sHsp action are the mode of substrate capture and the functional importance of sHsp oligomer dynamics. To address these, we engineered the structurally defined, dodecameric sHsps, Ta16.9 and Ps18.1, in order to restrict dissociation at each of two major oligomeric interfaces, dimeric and CT:ACD. A surprise on analysis of the proteins linked at the CT:ACD interface, Ta_CT-ACD_ and Ps_CT-ACD_, was that when oxidized the substructure of these sHsps proved to be covalent trimers, as opposed to covalent tetramers, as predicted from the available crystal structure (1GME) (17). This leads us to propose a tetrahedral structure for these dodecameric sHsps. Importantly, our analysis of the chaperone activity of Ta16.9 and Ps18.1 with fixed interfaces provides strong evidence that dissociation of the dodecamer at the CT:ACD interfaces is essential for optimal chaperone activity, and that stabilized dimers are more effective chaperones and likely to mediate the first encounter with denaturing substrate.

### A new geometry for the sHsp dodecamer

Our mutant studies, MS and modeling data support a new geometry for Ta16.9 and Ps18.1 dodecamers that is highly populated in solution. This dodecamer form has dimers arranged in a tetrahedral structure rather than in the double disk of the Ta16.9 crystal structure, and this tetrahedron is maintained in the disulfide mutants. The tetrahedral arrangement requires no changes in the sHsp monomer or dimer compared to the crystal structure, other than altering the angle of the CT relative to the ACD at a flexible hinge. In the double disk structure, the CT must adopt two different angles to form CT:ACD interfaces. In contrast, for tetrahedral geometry the CT in each monomer can have the same angle with respect to the ACD. Attempts to reduce and reoxidize Ta_CT-ACD_ under different conditions, including those used for crystallography, also only yielded covalent trimers. Thus, we suggest that the double disk geometry is a minor form in solution that is trapped by crystallization. The ability of sHsps to adopt different geometries accommodated by varying the angle of the CT while maintaining the CT-ACD interface intact is amply illustrated by different known sHsp oligomeric structures (2, 5, 52, 53). The ability to reassemble into different geometries, potentially using this same CT-ACD interface, may also contribute to the variable stoichiometries and morphologies of sHSP-substrate complexes (44, 54).

### sHsp temperature transitions and chaperone activity

Our thermal unfolding data indicate that stability of sHsp secondary structure is greatly enhanced when either of the dodecameric interfaces is linked by disulfides. Each Ta_CT-ACD_ and Ps_CT-ACD_ monomer has two interfaces with other monomers that are stabilized by disulfides, as opposed to Ta_dimer_, which has only one stabilized interface with its partner in the dimer. As a result, Ta_CT-ACD_ and Ps_CT-ACD_ are stabilized to a greater extent than Ta_dimer_. Unfolding profiles for different concentrations of the same protein will overlay when loss of secondary structure is independent of oligomeric state, while a shift to a higher melting temperature at higher concentrations indicates the protein is stabilized by association in a higher order structure. A cooperative transition in secondary structure is independent of concentration for the disulfide linked sHsps (Ta_CT-ACD_ and Ta_dimer_), implying that Ta_CT-ACD_ and Ta_dimer_ dissociate to well-folded covalent trimer and dimer respectively before unfolding. In contrast, unfolding of non-disulfide-linked sHsps occurs with deoligomerization. In summary, DLS studies show that the sHsp molecules undergo dissociation at both dimeric and CT-ACD interfaces upon heating. CD melt studies show that suboligomers (trimers and dimers) are folded at activity assay temperatures. Since stabilization of the dimeric interface enhances activity, and dissociation of CT-ACD interfaces is essential for activity, we infer that the folded dimers from sHsps are primary substrate capturing units.

### The two sHsps, Ta16.9 vs Ps18.1, differ in stability

Ta16.9 and Ps18.1 share 69% sequence identity (**Fig**. S1), but differ significantly in their ability to protect MDH, with Ps18.1 being more efficient (**Fig. 3**) (41). Previous molecular dynamics studies of Ps18.1 and Ta16.9 dimers suggest that Ps18.1 has larger exposed hydrophobic patches and that the Ps18.1 NT makes fewer contacts to itself compared to Ta16.9 (55). The surface area buried in the ACD dimers of Ta16.9 and Ps18.1 were calculated to be 2945 and 3059 Å^2^, using PDB files, 1GME (17) and 5DS2 (49), respectively. The somewhat larger buried surface area of the Ps18.1 dimer indicates a stronger dimeric interface, which would make it more stable than the Ta16.9 dimer. This is substantiated by the thermal melt data, from which the apparent Ta16.9 and Ps18.1 melting temperatures are estimated (from the first derivative plots at 10 "M) to be 48.3 and 50.6 ^o^C, respectively (**Fig. 2A**). The greater stability of Ps18.1 is also seen in DLS studies; Ps18.1 has a lesser tendency to form larger self-aggregates than Ta16.9, although both display dissociation to smaller species, including dimers. This difference in stability is likely a significant factor in the more efficient substrate protection by Ps18.1 compared to Ta16.9, and should be considered when assessing chaperone activity of other sHsps.

### sHsp dimers as the substrate capture unit

Our data support a model for sHsp activity as depicted in **Fig. 5**, in which the sHSP dimer makes the effective first encounter with denaturing substrate before assembly into higher mass sHsp-substrate complexes. By crosslinking the dimer interface in Ta_dimer_, we shifted the equilibrium of these dynamic dodecamers to the dimer form, while preventing dissociation at the CT:ACD interface essentially eliminated the dimer and monomer species (**Fig. 5A**). At the assay temperature of 45 ^o^C, all sHsps studied displayed dissociation to sub-oligomers; however, only suboligomers not linked at CT:ACD interfaces protected substrate. It is possible that linking the CT:ACD interface blocks substrate binding in the β4-β8 groove. However, our data, along with previous studies of Ta16.9 and Ps18.1 show that the NT is a major region involved in substrate (including MDH) binding (39, 41, 51). Studies of mammalian, yeast, archaeal and bacterial sHsps also have concluded that the NT is essential for substrate binding (56-59). Our data imply that the combined conformation of NT sites in the dimer are more effective in binding substrates than NT site conformations available in three linked monomers. Further, all the active sHsps retained complete (e.g. oxidized Ta_dimer_) or partial native-like secondary structure (e.g. Ta16.9, Ta_V144C_, reduced Ta_dimer_, Ps_V151C_) at 45 °C. Since the CD characteristics are primarily contributed by the ACD, and Ta_dimer_ is the most efficient chaperone (Fig. 5A), we infer that folded dimers are the primary substrate-capturing units. The recognition of substrates by oligomer dissociation products is an elegant mechanism of action, since each dodecamer comprises six dimers which diffuse much more rapidly. This allows for an efficient chaperone response, with subsequent assembly into larger complexes conferring stability over a longer life-time. The subunit dynamics and evidence that the NT of many sHsps is involved in substrate binding are consistent with our model being generally applicable to other sHsps interacting with diverse substrates.

**Figure 5:**
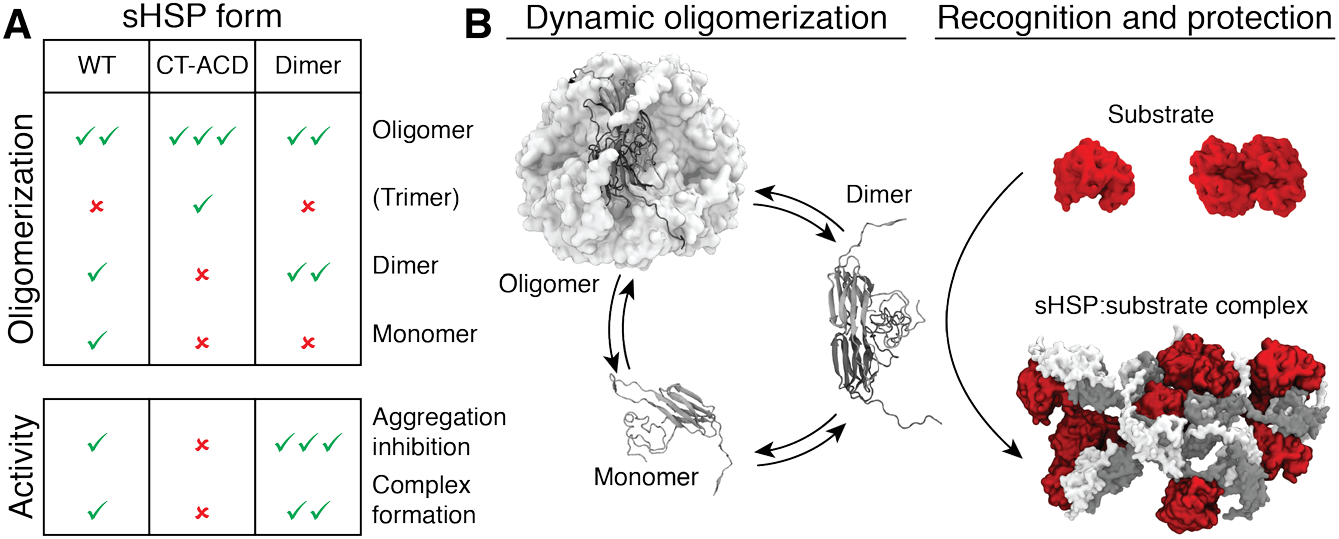
The sHsp dimer is the major substrate encounter species. **(A)** Table summarizing how the disulfide constrained sHsps (CT-ACD and Dimer) alter the population of dodecamer, dimer or monomer compared to wild type (WT) and resultant changes in chaperone activity. Check marks indicate the presence and abundance of a specific protein form or extent of chaperone activity, while crosses indicate absence of that protein form or activity. **(B)** Schematic showing how the dimer, which is the favored form in the dimer mutant and disfavored form in the CT-ACD mutants (as in A) is the major substrate capture form of the sHSP, followed by subsequent assembly of multiple dimers and substrates into heterogeneous complexes.

### Conclusions

Our data clearly support the importance of a specific mode of oligomer dissociation and the critical role of dimers in the mechanism of sHsp action. In addition to leading to a reassessment of the quaternary structure of these plant sHsps, analysis of the disulfide mutants shows that increasing stability of the dimeric interface enhances the sHsp holdase activity. Notably, it has also been observed that when stabilized with an interface disulfide, the core ACD of human αB-crystallin dimers was better at protecting substrates than wild type protein (60), and stabilizing crystallins has pharmacological potential in treating cataracts (61, 62). Thus, strategies to modulate the strength of the sHsp dimeric interface could be an approach to producing designer holdases active at different temperatures.

## MATERIALS AND METHODS

### Design and purification of disulfide mutants

The coordinates from the PDB file 1GME (17) for Ta16.9 were used for selecting positions to introduce cysteine residues for inter-polypeptide disulfide formation. The structure was analyzed using NACCESS (63) for identifying residues at the interaction surfaces. Pairs of residues on neighboring monomers, among which at least one residue was accessible (>7% accessibility) and were positioned such that Cα-Cα and Cβ-Cβ distances were <7.0 and <6.0 Å, respectively, were chosen (64, 65) to introduce disulfides at dimeric and non-partner interfaces (Table S1).

Ta16.9 and Ps18.1, had been previously cloned under the control of the IPTG-inducible T7 promoter (41). Cysteines were introduced by site-directed mutagenesis at the (i) dimeric interfaces: K49C W96C Ta16.9 and W48C H97C Ta16.9, and (ii) non-partner interfaces: E74C V144C Ta16.9 and V73C K145C Ta16.9. K49C W96C Ta16.9 and E74C V144C Ta16.9 were chosen for further study in comparison to Ta16.9 (**Fig. 1**). Single cysteine mutations: E74C Ta16.9, V144C Ta16.9, K49C Ta16.9, W96C Ta16.9 were also made. Corresponding mutations (K56C W103C Ps18.1, W55C R105C Ps18.1, D54C R105C Ps18.1, E81C V151C Ps18.1, E81C Ps18.1, V151C Ps18.1) were introduced in a homologue from *Pisum sativum*, Ps18.1, which shares 69% identity with Ta16.9 (Fig. S1). The residues in Ps18.1 for cysteine mutations were identified by sequence alignment of Ps18.1 and Ta16.9.

BL21-DE3 *E. coli* cells transformed with the required plasmid were grown in terrific broth at 37 °C to an O.D. of 0.6 before induction with 1 mM IPTG. Following induction, cells were grown at 25 °C for 12-16 hr. The cells were pelleted and then lysed by sonication in 20 mM Tris, pH 8.0, 20 mM NaCl. The supernatant following lysis and centrifugation (5000 x *g* rcf, 15 min), was loaded on a 40 mL Q Sepharose ion-exchange column and separated with a gradient of 20-250 mM NaCl of in 20 mM Tris, pH 8.0 spanning a total volume of 400 mL collecting 6 mL fractions. Fractions containing sHsp, which eluted between approximately 100-150 mM NaCl, were concentrated to 0.5 mL and separated on a S-200 Superdex GE gel filtration column equilibrated with 20 mM Tris, pH 8.0, 150 mM NaCl. All buffers used in purification contained 1 mM EDTA, 10 μM PMSF, 1 mM #-aminocaproic acid and 1 mM benzamidine. For purification of single cysteine sHsps, all buffers included 1 mM β-mercaptoethanol. Fractions containing purified protein were used for further analysis. Purity of the proteins and formation of disulfide bonds was ascertained by SDS-PAGE in the presence and absence of 10 mM DTT.

### Dithionitrobenzoic acid (DTNB) assay

The number of free cysteine residues in each of the proteins was determined by reaction with DTNB. Each of the proteins at 15 μM was incubated with 5 mM DTNB in 100 mM Tris, pH 8.0 and 4 M urea for 30 min at room temperature in the dark. 5 and 10 μM DTT samples were used as positive controls. Following incubation, absorbance of samples was measured at 412 nm to determine the concentration of thionitrobenzoic acid (TNB) using the extinction coefficient of 13,700 M^-1^ cm^-1^. The amount of TNB formed corresponds to the concentration of free thiols in the protein sample.

### (IM)-MS of Ta16.9, Ps18.1 and cysteine mutants

Proteins were prepared for native mass spectrometry at concentrations of 1-5 μM oligomer, in 200 mM ammonium acetate, pH 6.9. Mass spectra of all proteins were obtained on instruments (Q-ToF2, Synapt G1 HDMS; Waters Corp.) modified for the analysis of high mass species, according to methods described previously (66). Collision-induced dissociation was performed by acceleration into a dedicated collision cell within the instrument, either with or without selection of a particular ion population using the quadrupole analyzer (as specified). CCSs were obtained as described previously (67). Data were calibrated and analyzed using Masslynx software, and presented with minimal smoothing and the absence of background subtraction.

### Far UV CD spectroscopy

Protein samples for CD spectra collection were buffer-exchanged into 5 mM sodium phosphate, pH 7.5, 5 mM NaCl using 5 mL HiTrap desalting columns (GE Healthcare). The buffer included 0.5 mM TCEP as reducing agent for proteins Ta_V144C_, Ps_V151C_ and for reduced Ta_dimer_. Spectra were collected with 10 μM proteins on a Jasco J-1500 spectrometer. Spectra were collected at different temperatures using the temperature control program software. In the range of 25-85 ^o^C, spectra were collected at data intervals of 10 °C. The rate for temperature increase was set to 2 °C min^-1^, with the temperature held constant during data collection. The scan rate, data pitch, response time and bandwidth parameters were set to 100 nm min^-1^, 1 nm, 2 sec and 1 nm, respectively. An average of three spectra collected for each sample in the range of 250-200 nm are reported after subtraction of corresponding buffer spectra and smoothing. All samples after data collection were incubated at room temperature for 15 min, and then spectra were collected with the parameters as above to check for protein refolding.

### Thermal stability of sHsp secondary structure

Thermal stability of sHsp secondary structure was assessed on a Jasco J-1500 CD spectrometer by monitoring sample ellipticity at 205 nm as a function of temperature. Measurements were made for 1.0 and 10.0 μM protein in 5 mM sodium phosphate, pH 7.5, 5 mM NaCl, in 1 cm and 1 mm path length cuvettes, respectively. A scan rate of 1 °C min^-1^ with a data pitch of 0.5 °C was used.

### Dynamic light scattering (DLS)

DLS profiles were collected for protein samples in 25 mM HEPES, pH 7.5, 150 mM KCL, 5 mM MgCl_2_ on a Malvern Zetasizer Nano instrument. For reduced protein samples the buffer included 20 mM DTT. Data were collected for 50 μL, 10 μM protein samples in a micro-quartz cuvette (Malvern-ZEN2112). Protein samples were filtered through a 0.22 μm filter prior to measurement. Each sample was incubated at 25 °C for 10 min in the cuvette in the instrument set to 25 °C. Following incubation, three measurements were made, each consisting of 15 runs of 15 sec each. The cuvette holder temperature was then increased by 10 ^°^C. The same sample was then incubated at the increased temperature for 10 min prior to data collection as before. This was repeated until data collection at 65 °C was complete. The sample was then cooled to 25 °C, incubated for 10 min, and DLS measurements repeated. Data were analyzed using the software Zetasizer 7.11. Corrections for viscosity and refractive index of water with respect to temperature were taken into account for calculating size distributions. The derived count rates were used for generating scatter intensity plots as a function of temperature.

### Activity assays for protection of substrate protein by sHsps

The substrate protein used was porcine heart malate dehydrogenase (MDH). All protein concentrations are cited for protein monomers. 3 "M MDH was incubated with different concentrations of sHsp in 25 mM HEPES, pH 7.5, 150 mM KCl, 5 mM MgCl_2_. For assays of sHsps in the reduced state, 20 mM DTT was included in the buffer. Reaction mixtures were incubated at 45 °C for 2 h, and then cooled on ice for 10 min, as described previously (41). To ascertain the extent of MDH precipitation, reaction mixtures were diluted 8-fold with assay buffer, and sample scattering intensity was measured on a Photon Technology International fluorometer. Excitation and emission wavelengths were set to 500 nm with both slit widths set to 2 nm. Scattering intensity was measured over 20 sec, with a data pitch of 1 sec. Measurements were averaged over the 20 sec period. For analysis of sHsp-MDH complex formation by analytical SEC, 20 μ/ of supernatant obtained after centrifugation (20 min at 16,000 x g) for each reaction mixture was injected on a TSKgel Ultra SW aggregate column (Tosoh). The column was run at 1 mL^-1^ in 5 mM sodium phosphate pH 7.5, 150 mM KCl, 5 mM MgCl_2_, and absorbance at 220 nm of the eluent was recorded.

### SAXS

Data were collected at the B21 bending-magnet instrument at the Diamond Light Source (Harwell, UK). Samples were prepared in 200 mM ammonium acetate to a concentration of 5 mg ml^-1^ and two successive 2-fold dilutions. Protein and corresponding buffer solutions were exposed to the beam in a 1.6 mm diameter quartz capillary at 15 °C. The sample capillary was held in vacuum, and subjected to a cleaning cycle between each measurement. A Pilatus 2M two-dimensional detector was used to collect 180-frame exposures of 1 s from each sample and the corresponding buffer. The detector was placed at 3.9 m from the sample, giving a useful Q-range from 0.012 Å^-1^ to 0.4 Å^-1^. Two-dimensional data reduction consisted of normalization for beam current and sample transmission, radial sector integration, background buffer subtraction and averaging. Each frame was inspected manually and discarded if signs of radiation damage were apparent. Data scaling, merging and Guinier analysis were performed in PRIMUS (68).

## ACKNOWLDGEMENTS

The authors thank Dr. Lizz Bartlett for her assistance with the UMass Amherst Institute for Applied Life Sciences Core Facility for Biophysical Characterization, Dr. Eugenia Clerico for useful discussions, and Umaru Barrie for help with preparation of the single cysteine mutants. We acknowledge access to B21 and help from Mark Tully and James Doutch at the Diamond Synchrotron (SM9384-2 to JLPB). The work was supported by the Engineering and Physical Sciences Research Council (EP/J01835X/1 to JLPB, EP/P016499/1 to MTD), and the United States National Institutes of Health RO1 GM42762 (to EV).

